# Matters arising: Forced Symmetry Artifacts in a Prohibitin Complex Structure

**DOI:** 10.1101/2025.05.16.653159

**Authors:** Eric Herrmann, Kevin M. Rose, James H. Hurley

## Abstract

In a recent publication, Lange et al.^1^ proposed an 11-spoked wheel model for the mitochondrial prohibitin complex consisting of six PHB1 and five PHB2 subunits built into low resolution density obtained from 11-fold symmetry averaging. The proposed unequal stoichiometry is inherently incompatible with symmetry. The building of an asymmetric molecular model into density obtained by 11-fold symmetry averaging is inherently self-contradictory. This contradiction alone calls the validity of the derived structure into question. Our own data revealed that the complex primarily adopts an asymmetric open conformation in situ. We show that it was the incorrect symmetry imposition on the open conformation that led to the spoked wheel density reported by Lange et al.. Further validation by re-analysis of crosslinking-mass spectrometry data, and a test of in situ template matching all support a larger prohibitin complex architecture. These findings underscore the need for great care in the imposition of symmetry cryo-ET at less than atomistic resolution.

---

In a recent publication, Lange et al.^1^ proposed an 11-spoked wheel model for the mitochondrial prohibitin complex (composed of PHB1 and PHB2 subunits) based on *in situ* cryo-ET data. They described a spoke-like assembly of 11 subunits, six PHB1 and five PHB2 subunits built into a density reconstruction obtained from 11-fold symmetry averaging. While PHB1 and PHB2 share similarity, the proposed unequal stoichiometry is inherently incompatible with symmetry. The building of an asymmetric molecular model into density obtained by 11-fold symmetry averaging is inherently self-contradictory. This contradiction alone calls the validity of the derived structure into question.

Our laboratory has independently obtained an in situ cryo-ET reconstruction of human prohibitin^2^. Our data revealed two major structural states: a near-symmetric, closed dome-shaped conformation (Fig. 1a) and an open, C-shaped conformation in which several subunits are flexible and unresolved (Fig. 1b). To test the hypothesis that enforcing symmetry at an incorrect stage in processing led to an unphysical spoke-like density map, we reanalyzed our own cryo-ET data for the open and closed states, and for a merged particle stack combining both populations (Fig. 1c). All three sets were subjected to C11 symmetry enforcement. The resulting reconstructions showed dramatic differences: the closed population produced a dome-shaped density (Fig. 1d), while the open population yielded a spoke-like morphology (Fig. 1e) similar to that reported by Lange et al. The merged dataset displayed a hybrid (Fig. 1f). Our analysis of mitochondrial samples indicated that under healthy conditions, approximately 75% of prohibitin complexes are in the open state and only 25% are closed. Thus, symmetry imposition on a predominantly open, asymmetric population artificially generates a spoke-like reconstruction. While applying C11 symmetry led to a minor improvement in apparent resolution, pathological features in the Fourier Shell Correlation (FSC) curves strongly suggest that true symmetry was absent. Normally, the FSC curve should taper off smoothly to zero as resolution increases, reflecting diminishing correlation at high spatial frequencies. Instead, we observed multiple sharp and unnatural peaks. These jumps imply that the reconstruction is echoing symmetry-related noise, not genuine signal. This highlights a broader concern: in flexible, heterogeneous datasets, enforcing symmetry can average out conformational diversity and lead to misleading structural models.

**Figure 1:**
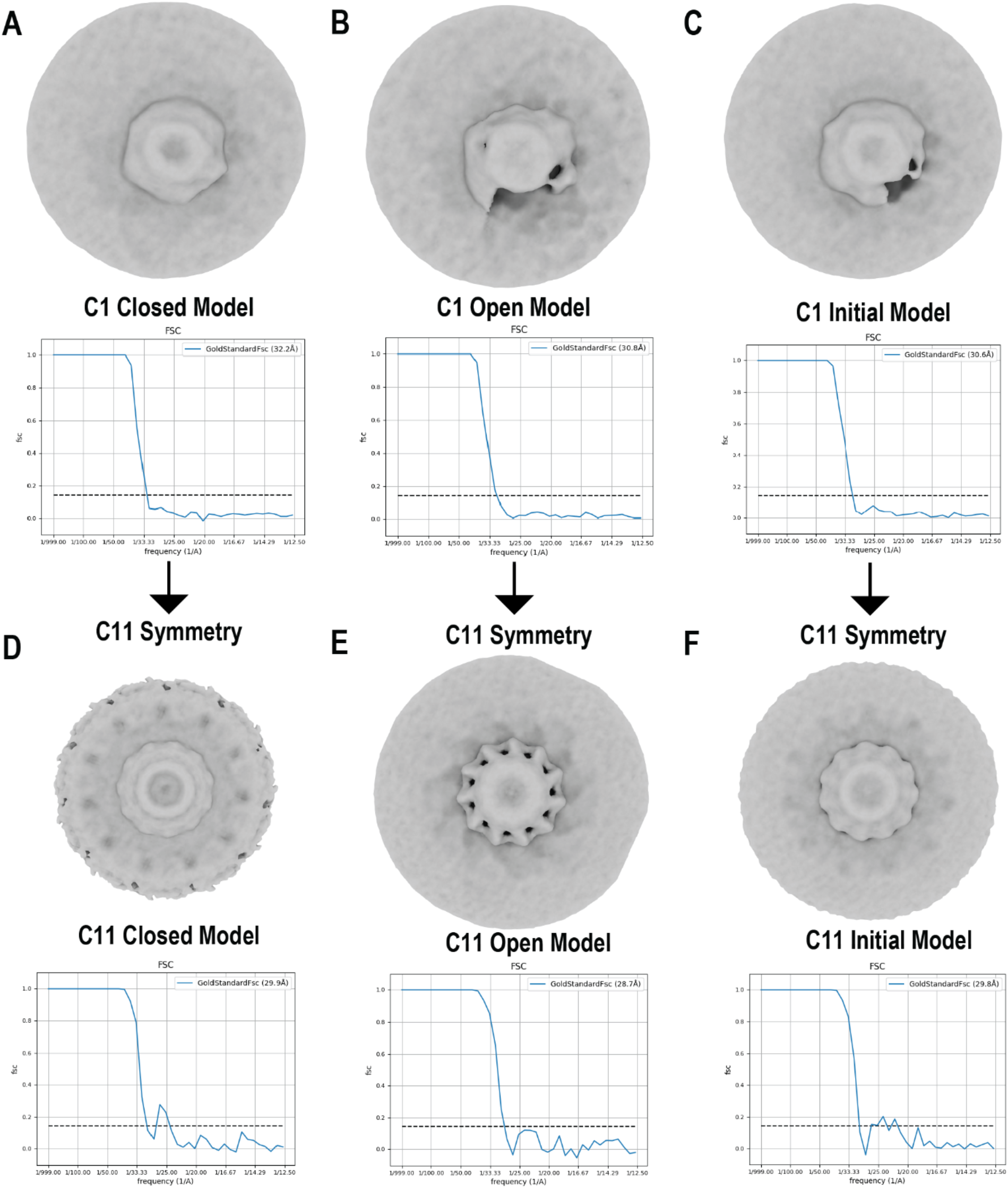
Effects of symmetry enforcement on prohibitin conformations.(A–C) Cryo-ET reconstructions of the prohibitin complex in its closed, dome-shaped state (A), open C-shaped state (B), and a combined dataset of both states (C), prior to symmetry application.(D–F) Reconstructions after applying 11-fold (C11) symmetry to the closed (D), open (E), and combined (F) particle populations, with corresponding Fourier Shell Correlation (FSC) curves shown. Symmetry imposition on the flexible open state produces an artificial spoke-like morphology. FSC curves show multiple sharp peaks, indicating artifacts introduced by symmetry enforcement, rather than genuine high-resolution signal.

Further analysis of the molecular architecture supports our alternative model. Our reconstructed prohibitin complex, consisting of twelve PHB1–PHB2 heterodimers (approximately double the mass of the Lange et al. model), fits the established architecture of SPFH domain-containing protein families, such as flotillins^3^ and the bacterial HflK/C complex^4^. Strikingly, crosslinking mass spectrometry (XL-MS) data, also used by Lange et al. for validation^5^ show improved match rates and distance satisfaction with our model, further calling their interpretation into question (Figs. 2a–c). To further validate the structural fit, we used PyTom^6^ with a 30 Å dilated mask of both models to 3D template match prohibitin within our tomograms (Fig. 2d). The local cross-correlation scores of individual particle picks were consistently higher for our model compared to the model proposed by Lange et al. (Fig. 2e), indicating closer agreement with the experimental data.

**Figure 2:**
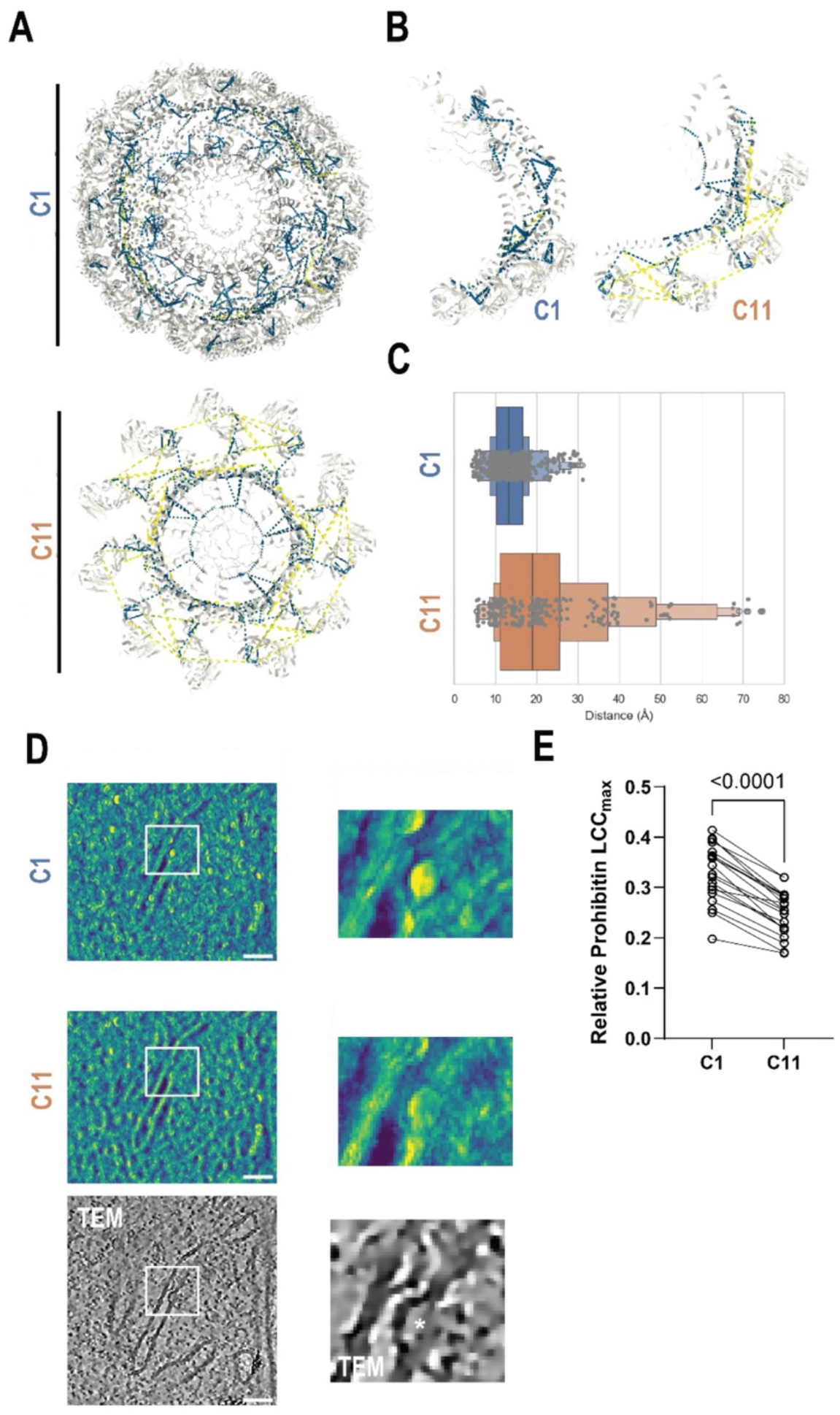
Validation of the prohibitin model by crosslinking mass spectrometry and template matching. (A) Mapping of crosslinking mass spectrometry (XL-MS) restraints onto our reconstructed prohibitin model (top) compared to the Lange et al. model (bottom) colored by distance (<24Å = blue, >24Å = yellow).(B) Close-up view highlighting crosslink satisfaction between two adjacent heterodimers. (C) Quantitative plot of crosslink distances, showing a higher proportion of satisfied restraints (shorter distances) in our model.(D) Local cross-correlation maximum (LCCmax) scoring results from 3D template matching using PyTom, next to a denoised tomogram (scale bar = 100 nm). Higher scores (yellow) indicate better fit to experimental data. (E) Scatter plot of individual particle LCCmax scores, demonstrating a statistically significant improvement in template matching for our model compared to the Lange et al. model.

While writing this article, two preprints appeared describing the single-particle cryo-EM structure of a detergent-extracted *Chaetomium thermophilum*^7^ *and homo sapiens*^8^ prohibitin complex at ∼3.5 Å resolution. In that study, PHB1 and PHB2 subunits were clearly resolved in an organization closely resembling our model, but with 22 instead of our proposed 24 subunits. Due to the limited resolution of our *in situ* cryo-ET data, we were unable to precisely determine the subunit count and relied on structure prediction tools. Nevertheless, these findings agree with our data and challenge the spoke-like model proposed by Lange et al.

Taken together, our findings highlight the importance of applying symmetry constraints cautiously, especially in the context of flexible and dynamic structures. Extensive validation — as demonstrated in recent flotillin complex studies^9^, and, notably, in the *C. thermophilum* prohibitin preprint where symmetry was only applied after reaching ∼8 Å resolution^7^ — is essential to distinguish true biological architecture from reconstruction artifacts. As cryo-EM and cryo-ET increasingly tackle heterogeneous *in situ* assemblies, careful symmetry assessment will be critical to maintain the reliability of structural models.

## EMDB and PDB accession codes

The spoke-like prohibitin from Lange et al. can be found with accession codes EMD-19459 and PDB:8RRH. Our dome-like prohibitin can be found with accession codes EMD-70179 and PDB:9O6S in the closed state and EMD-70180 and PDB:9O6T in the open state. The codes for the high-resolution structure of the *C. thermophilum* prohibitin are not released at the time this manuscript was written, but the *homo sapiens* prohibitin can be found under EMD-70267 and PDB: 9O9Z.

## Materials and Methods

### Cryo-electron tomography data processing

A detailed description of data collection and tomogram reconstruction is available in our corresponding publication^2^. Briefly, 2,677 particles were manually picked from 76 tilt series and processed using RELION 5.0^10^. After 3D classification, 1,193 particles were assigned to the closed class, 1,156 to the open class, and 328 were discarded as trash. Particles from the open class, closed class, and a combined set were refined using C1 or C11 symmetry. The resulting volumes and Fourier Shell Correlation (FSC) curves were used for visualization.

### Crosslink data analysis

Crosslinking data^5^ were analyzed using the XMAS^11^ bundle in ChimeraX^12^. A distance cutoff of 24 Å was applied to define close matches.

### Template matching

Template matching was performed in PyTOM^6^ using a 30 Å dilated mask of our model and the model proposed by Lange et al.^1^ on our tomograms with a pixel size of 10.5 Å/px. The local cross-correlation scores were imported into Fiji^13^ for further analysis using the line scan tool. The maximum score per particle was used for paired Student’s t-test analysis in GraphPad Prism and visualized as a scatter plot.

## Acknowledgments

This research was funded by Aligning Science Across Parkinson’s [ASAP-000350] through the Michael J. Fox Foundation for Parkinson’s Research (MJFF) (J.H.H.) and by the Alexander von Humboldt Foundation (E.H.).

